# Social and asocial learning in zebrafish are encoded by a shared brain network that is differentially modulated by local activation

**DOI:** 10.1101/2021.10.12.464038

**Authors:** Júlia Pinho, Vincent Cunliffe, Giovanni Petri, Rui F. Oliveira

## Abstract

Group living animals can use social and asocial cues to predict the presence of a reward or a punishment in the environment through associative learning. The degree to which social and asocial learning share the same mechanisms is still a matter of debate, and, so far, studies investigating the neuronal basis of these two types of learning are scarce and have been restricted to primates, including humans, and rodents. Here we have used a Pavlovian fear conditioning paradigm in which a social (fish image) or an asocial (circle image) conditioned stimulus (CS) have been paired with an unconditioned stimulus (US=food), and we have used the expression of the immediate early gene *c-fos* to map the neural circuits associated with social and asocial learning. Our results show that the learning performance is similar with social (fish image) and asocial (circle image) CSs. However, the brain regions involved in each learning type are distinct. Social learning is associated with an increased expression of *c-fos* in olfactory bulbs, ventral zone of ventral telencephalic area, ventral habenula and ventromedial thalamus, whereas asocial learning is associated with a decreased expression of *c-fos* in dorsal habenula and anterior tubercular nucleus. Using egonetworks, we further show that each learning type has an associated pattern of functional connectivity across brain regions. Moreover, a community analysis of the network data reveals four segregated functional submodules, which seem to be associated with different cognitive functions involved in the learning tasks: a generalized attention module, a visual response module, a social stimulus integration module and a learning module. Together, these results suggest that, although there are localized differences in brain activity between social and asocial learning, the two learning types share a common learning module and social learning also recruits a specific social stimulus integration module. Therefore, our results support the occurrence of a common general-purpose learning module, that is differentially modulated by localized activation in social and asocial learning.

## Introduction

The social intelligence hypothesis (1,2) states that living in social groups creates a demand for enhanced cognitive abilities in order to handle the variability and unpredictability of social interactions, hence driving the evolution of more complex cognitive skills (aka intelligence), and consequently selecting for larger executive brains (i.e. social brain hypothesis) (3,4). However, two different scenarios have been proposed for how these evolved cognitive abilities implement adaptive behavior. According to a general-purpose brain scenario, mechanisms of information input, encoding, storage and retrieval are shared between functional domains (e.g. social, foraging, predator avoidance), hence, although evolved in a specific domain (e.g. social), enhanced cognitive abilities are advantageous in all domains. Alternatively, each functional domain relies on special-purpose cognitive modules, which are highly specialized with independent mechanisms of information processing. In this regard, there is an ongoing debate in the field of social cognition, on the extent to which social learning (i.e. learning from other individuals) is a general-domain or a domain-specific process (5–9). For example, comparative studies in birds and primates show correlations between the performance on social learning and individual learning (aka asocial learning) tasks or measures of behavioral flexibility, suggesting that these traits evolved together (10–12). Furthermore, observational learning in bumblebees has been shown to emerge through the integration of two learned associations following Pavlovian conditioning rules (13). In contrast, there is also comparative evidence supporting the occurrence of domain-specific modules, such as the differences found in social learning, but not in individual learning, between two corvid species with differences in degree of sociality, or between human children and apes (14,15). Moreover, intra-specific studies in mice show that maternal deprivation early in life impairs social learning whereas spatial learning is unaffected, and that communally-reared mice, when compared to single-mother reared mice, have better social competence but do not differ in spatial learning and memory capacity tests (16,17). Finally, a third scenario has also been proposed that suggests that social learning operates on the same general learning mechanisms as asocial learning with adaptive specializations present only for the input systems (i.e. social information acquisition) (6–9).

The study of the proximate mechanisms (i.e. genetic basis, neural circuits) of social and asocial learning can, in principle, help to clarify the occurrence of shared processes. Unfortunately, there are few studies on such mechanisms with notable exceptions for the study of observational fear learning and for social learning of food preference. In both humans and rodents, social (i.e. observational) and asocial fear conditioning share, at least partially, the same neural substrates, with the anterior cingulate cortex (ACC) processing the social information and conveying it to the amygdala, which plays then a major role in the CS-US pairing in both learning types (18–22). In rodents, a specialized olfactory subsystem has been described that is required for the acquisition of socially transmitted food preferences (23). Moreover, social fear learning and classic fear learning can prime each other (i.e. a prior observational fear learning will enhance fear conditioning and vice-versa). Together these results suggest an overlap of the neural mechanisms involved in social learning and in learning from direct experience, with specializations being mainly present at the level of social information acquisition. However, the above-mentioned studies address specific brain regions that are chosen a priori as candidates for the social learning tasks, and studies using a unbiased brain network approach are lacking in this field. This is particularly important because despite the overlap of brain circuits processing social and asocial learning, the candidate brain region approach does not rule out the occurrence of specialized circuits elsewhere in the brain. Furthermore, the analysis of localized neuronal activation, which is usually the parameter studied in relation to the behavioral output, does not provide per se information on the patterns of co-activation across a brain network that may reveal either specialized or conserved modules for the two learning types. Finally, the study of the neural mechanisms of social learning has focused on mammals, and comparative data in other vertebrate species that lack evolved cortical structures is also missing.

Here we have used a classic (Pavlovian) conditioning paradigm in zebrafish, in which a social (image of a zebrafish) or an asocial (image of a circle) conditioned stimulus (CS) was paired with an unconditioned stimulus (US = food), in order to investigate the neural basis of social and asocial learning. The choice of this classic conditioning paradigm where the social and asocial treatments are matched for everything except the visual shape of the CS rule out putative confounding variables, such as for example the involvement of different sensory modalities in the acquisition of the social information. We have used the expression of the immediate early gene *c-fos* as a molecular marker of neuronal activity (24,25). We have also developed a method to analyze brain network functional connectivity based on the brain regions’ co-activation matrices for each experimental treatment. In network neuroscience, such matrices are built for individual brains based on a similarity measure between the timeseries of brain parcels in fMRI, or the different channels in an EEG (26). In the case of a molecular marker of neuronal activity, such as *c-fos*, that only provides a single snapshot per brain, the correlations of activity between brain regions (i.e. the number of positive *c-fos* cells for each pair of brain nuclei), is obtained for a set of brains from different individuals. Therefore, the estimation of the similarity in activation between regions can only be computed at the group level (i.e. using datapoints from individuals in the same treatment), effectively extracting a group-level, as opposed to individual, functional connectivity network. Finally, we used techniques from network community analysis to identify coherent sets of nodes (submodules) consistently recruited by the different cognitive tasks involved in social and asocial learning.

## Results

### Social and asocial classic conditioning in zebrafish

Pavlovian conditioning was assessed using a plus-maze paradigm divided into a training phase and a probe test. During the training phase, we spatially paired a social or an asocial conditioned stimulus (CS) with an unconditioned stimulus (US; food = bloodworms) in a specific location (Fig. 1a). The percentage of right choices per session (composed of 8 trials) was measured. In the probe test (24h after the last training session), individuals only had access to the CS, and the time spent in the region of interest (RoI) of the correct arm of the maze was quantified to measure learning. Unpaired treatments were used as controls, where the CS (either social or asocial) was spatially unmatched with the US.

**Figure 1.**
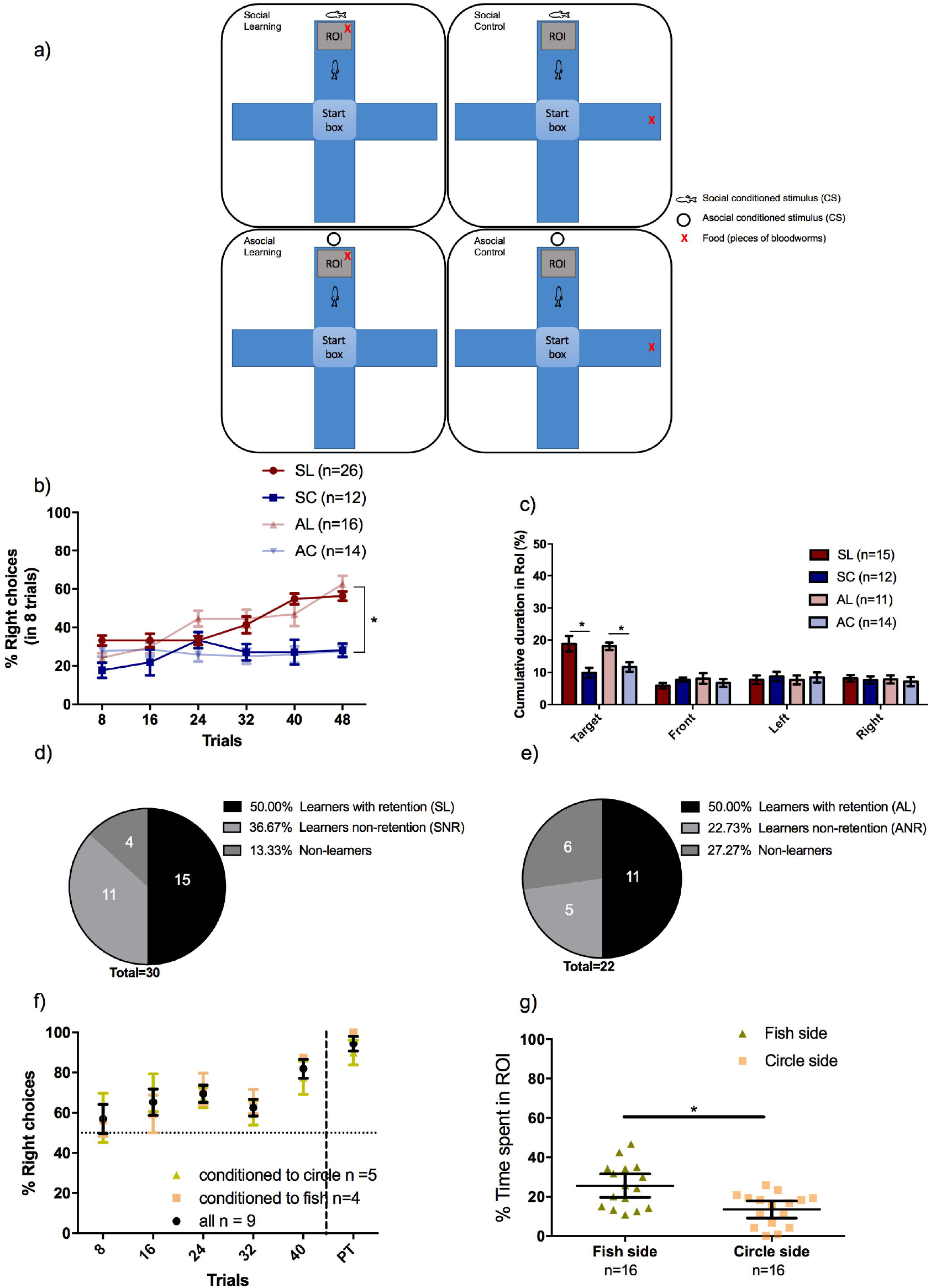
Social and asocial classic conditioning in zebrafish. (A) Schematic representation of the plus-maze paradigm: 4 groups observed a CS (social or asocial cue) paired with a US (food: bloodworms) in the same arm (paired treatments: SL and AL) hence being able to establish the CS-US association; or in different arms (unpaired treatments: SC and AC) the controls of the experiment. (B) During the training phase animals increased significantly the percentage of right choices both in the social learning (SL, in red circles) and asocial learning (AL, in light red circles) treatments in comparison with the respective unpaired treatments [in blue circles social unpaired control (SC) and in light blue circles the asocial unpaired control (AC)]. (C) In the probe test, the cumulative duration of time spent in the RoI indicates that learners (social and asocial) increased the time spent in the target arm. Pie graphs indicate the proportion of learners, non-learners and non-retention animals in social (D) and asocial conditions (E). The ability of the animals to distinguish between the social and asocial stimuli used in this experiment was tested by conditioning the animals to approach one stimulus and avoid other, independent of their initial preference (in yellow triangles animals conditioned to approach asocial, in light pink squares individuals conditioned to approach social and in black circles the average of all individuals) (F).The preference for the social and asocial stimuli [fish (yellow circle) or circle (grey square) static 2D picture, respectively] was assessed using a preference test (G). Asterisks indicate statistical significance at p < 0.05 using planned comparisons.

Animals learned both socially and asocially (learning main effect: X^2^_R_(1)=28.44, p<0.0001) as shown by the comparison in the percentage of correct choices between paired CS-US [social learning (SL) and asocial learning (AL)] and unpaired CS-US [social control (SC) and asocial control (AC)) treatments for social and asocial CSs (SL vs SC:X^2^_R_(1)=6.95, p=0.0089; AL vs AC:(X^2^_R_(1)=28.44, p<0.0001) (Fig. 1b). Animals in the social and asocial learning treatments (SL and AL)) acquired information at the same rate, since no significant differences between social and asocial learning curves were found either in slope (X^2^_R_(1)=1.53, p=0.22) or elevation (X^2^_R_(1)=0.001, p=0.97) (Fig. 1b). It is worth mentioning that there were also no significant differences between the social and asocial control treatments either in slope (X^2^_R_(1)=2.0, p=0.16) or elevation (X^2^_R_(1)=0.14, p=0.70) (Fig. 1b). Moreover, in control treatments, animals did not present any biased-behavior towards one of the arms of the plus maze exhibiting a random proportion of choices over the trials (25% in social and asocial treatments across the training sessions).

In the probe test, individuals from the learning treatments (SL and AL) spent more time in the target arm independently if they were trained using a social (X^2^_F_(1)=12.89, p=0.001) or an asocial (X^2^_F_(1)=11.53, p=0.001) CS, when compared to the control treatments (i.e. unpaired CS-US). We did not observe any significant difference in the time spent in the other arms of the plus-maze indicating an absence of any spatial biases in the spatial use of the maze by the fish during this phase of the experiment (opposite arm to the target arm: SL vs SC,, X^2^_F_(1)=1.22, p=0.276, AL vs AC,, X^2^_F_(1)=0.32, p=0.577; left of the target arm: SL vs SC, X^2^_F_(1)=0.42, p=0.522, AL vs AC, X^2^_F_(1)=0.06, p=0.801; right of the target arm: SL vs SC, X^2^_F_(1)=0.25, p=0.617, AL vs AC, X^2^_F_(1)=0.21, p=0.649) (Fig. 1c).

This paradigm allowed the classification of individuals in the learning treatments (SL and AL) into three different categories: non-learners, learners and learners that forget the learned information from the last learning session to the probe test (i.e. no-retention group: social non-retention (SNR) and asocial non-retention (ANR)).The learners were able to acquire the information and recall it (50% of individuals in both the social and asocial learning treatments); the “no-retention group” were animals that despite showing a learning curve during the training sessions did not recall the acquired information in the probe test (36.67% individuals in the social group (SNR) and 22.73% individuals in the asocial group(ANR)); and a small percentage of individuals that did not improve the performance over the training sessions (13.33% individuals in the social groups and 27.27% individuals in the asocial group) were classified as non-learners (Figs. 1d, e). The non-learners and no-retention animals were identified using the interval of confidence in the training phase and duration in the ROI of the target arm during the probe test as criteria, respectively. The proportion of learners (X^2^(1)=0, p=1), non-learners (X^2^(1)=1.56, p=0.21) and non-retention (X^2^(1)=1.14, p=0.29) individuals did not differ between social and asocial learning treatments (Figs. 1d, e).

Given the lack of difference in behavioral measures between social and asocial learning it was important to make sure that the individuals can discriminate the two CS stimuli used in this test. Thus, a visual discrimination task to asses if zebrafish can discriminate between the two stimuli (social and asocial CS) was used, where one stimulus was associated with a reward and the other with a punishment (e.g. social stimuli as a reward and asocial as a threat, and vice-versa). This test indicated that zebrafish could distinguish between the social and asocial stimuli used in this study and that the learning curve for the acquisition of these discrimination was similar when either the social or the asocial were paired with the reward (slope X^2^_R_(1)=1.74, p=0.22; elevation (X^2^_R_(1)=0.43, p=0.53; Fig. 1f). Given that social animals usually have an innate preference for social cues we have also tested the preference of zebrafish for the social stimuli used here to make sure that it had a positive valence. Preference was assessed using a choice test, where animals could choose between spending time near the social vs. the asocial stimuli used in our study. As predicted, a preference for the social stimulus was observed (t(15)=2.55, p=0.02; Fig. 1g).

In summary, in zebrafish both social and asocial cues are equally efficient as a CS in a classic conditioning paradigm, despite zebrafish having an innate preference for the social cue.

### Brain regions associated with social and asocial classic conditioning in zebrafish

The brain regions (see Table 1 for list of regions studied and their abbreviations) associated with social and asocial learning were determined using the expression of the immediate early gene *c-fos*, a marker of neuronal activation, by *in situ* hybridization. Because of possible laterality effects the expression of *c-fos* was measured on both brain hemispheres (noted below as left or right for each brain region). We identified as brain nuclei involved in social (SL) or asocial (AL) learning those that presented significant differences in *c-fos* positive cells between animals of the paired treatments (SL or AL) that were able to acquire and recall the CS and the respective unpaired control treatments (SC or AC, respectively). The following areas showed increased activation associated with social learning: olfactory bulb (OB) (left: X^2^_F_(1)=8.87, p=0.022; right: X^2^_F_(1)=7.35, p=0.022), ventral nucleus of ventral telencephalic area (vV) (left: X^2^_F_(1)=6.42, p=0.048; right: X^2^_F_(1)=5.72, p=0.048), ventral habenular nucleus (Hav) (left: X^2^_F_(1)=6.06, p=0.04; right: X^2^_F_(1)=10.28, p=0.012) and ventral medial thalamic nucleus (VM) (left: X^2^_F_(1)=6.20, p=0.038; right: X^2^_F_(1)=7.46, p=0.011) (Table 2; Figs. 2a-h). On the other hand, the left dorsal habenular nucleus (Had_l_) and the right anterior tubercular nucleus (ATN_r_) were differentially activated during asocial learning (X^2^_F_(1)=6.86, p=0.05 and X^2^_F_(1)=8.42, p=0.028, respectively; Table 2; Figs. 2i-l).

**Table 1.** Nomenclature of brain regions and their list of abbreviations used in the present work. The letters l and r were added as subscripts to identify the left and right hemispheres.

| Brain regions | Abbreviations |
| --- | --- |
| Olfactory bulbs | OB |
| Dorsal Telencephalic area | D |
| Dorsal nucleus of ventral telencephalic area | Vd |
| Central nucleus of ventral telencephalic area | Vc |
| Ventral nucleus of ventral telencephalic area | Vv |
| Lateral nucleus of ventral telencephalic area | VI |
| Central zone of dorsal telencephalic area | Dc |
| Lateral zone of dorsal telencephalic area | DI |
| Medial zone of dorsal telencephalic area | Dm |
| Posterior zone of dorsal telencephalic area | Dp |
| Supracommissural nucleus of ventral telencephalic area | Vs |
| Dorsal zone of dorsal telencephalic area | Dd |
| Anterior part of parvocellular preoptic nucleus | Ppa |
| Postcommissural nucleus of ventral telencephalic area | Vp |
| Magnocellular preoptic nucleus | PM |
| Posterior part of parvocellular preoptic nucleus | PPp |
| Dorsal habenular nucleus | Had |
| Ventral habenular nucleus | Hav |
| Anterior thalamic nucleus | A |
| Ventromedial thalamic nucleus | VM |
| Ventrolateral thalamic nucleus | VL |
| Ventral zone of periventricular hypothalamus | Hv |
| Anterior tuberal nucleus | ATN |
| Lateral hypothalamic nucleus | LH |
| Dorsal zone of periventricular hypothalamus | Hd |
| Central posterior thalamic nucleus | CP |
| Periventricular nucleus of posterior tuberculum | TPp |
| Periventricular gray zone of optic tectum | PGZ |
| Caudal zone of periventricular hypothalamus | Hc |
| Diffuse nucleus of the inferior lobe | DIL |
| Dorsal tegmental nucleus | DTN |
| Central nucleus of the inferior lobe | CIL |
| Nucleus lateralis valvulae | NLV |
| Griseum central | GC |
| Corpus mamillare | CM |

**Table 2.**
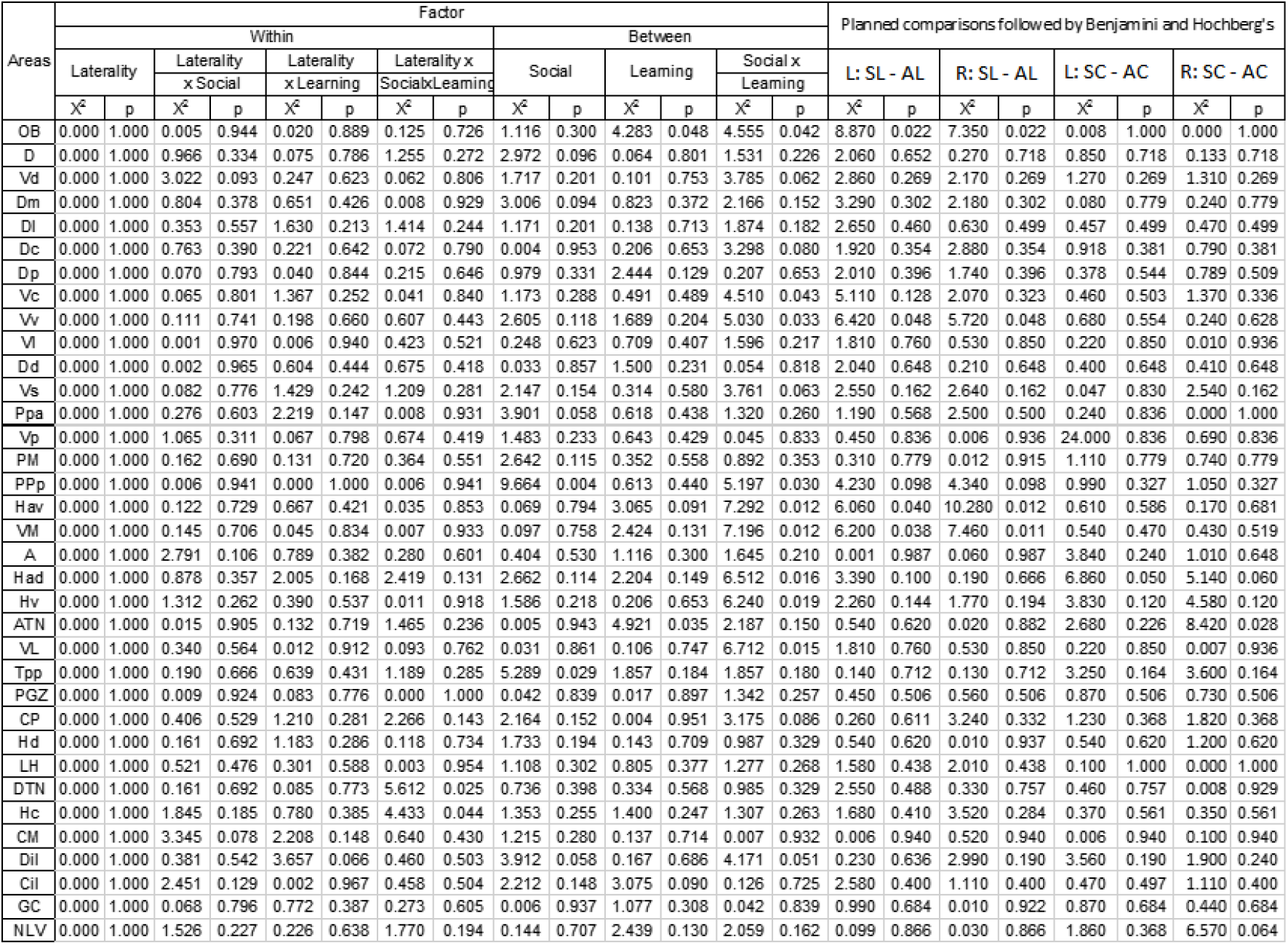
Effect of social and asocial learning on neuronal activation in different brain nuclei assessed by quantification of *c-fos* positive cells. Main effects of learning (learning vs. non-learning) and social context (i.e. social vs. asocial), interaction (learning x social context) and multiple comparisons were computed using non-parametric Friedman Test. SL, social learners (SL); SC, social controls; AL, asocial learners; AC, asocial controls. See list for abbreviations.

**Figure 2.**
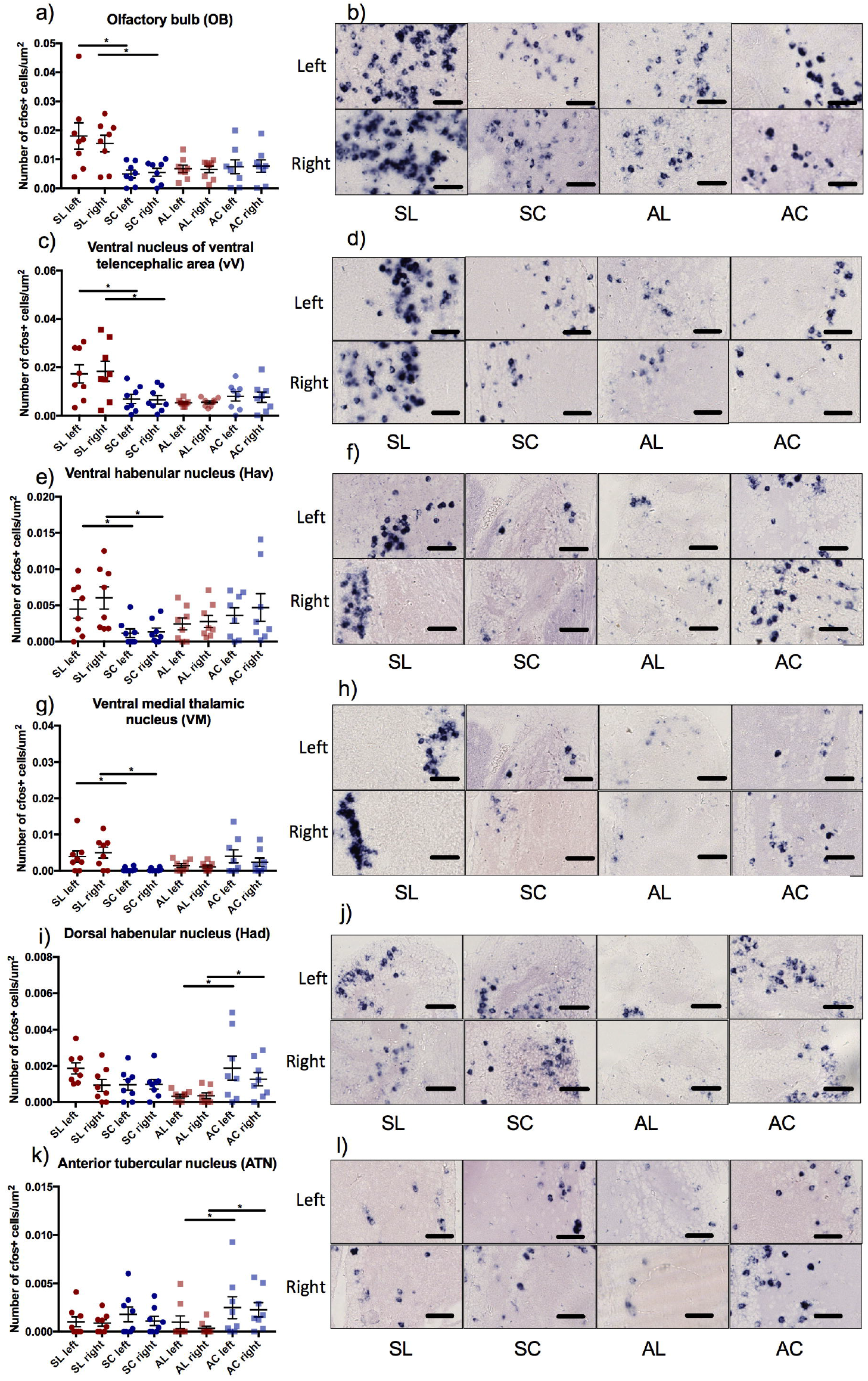
Neuronal activity associated with social (A – H) and asocial (I – J) classic conditioning in zebrafish assessed by *in situ* hybridization of the immediate early gene *c-fos*. Representative photomicrographs of *c-fos* in situ hybridization in areas that present significant differences associated with learning: OB (B), Vv (D), Hav (F), VM (H), Had (I) and ATN (K). Asterisks indicate statistical significance at p < 0.05 using planned comparisons followed by Benjamini and Hochberg’s method for multiple comparisons p-value adjustment. Scale bars represent 40 μm. See Table 1 for abbreviations of brain regions.

In summary, despite the behavioral similarities between social and asocial learning in zebrafish described in the previous section, learning from a social CS is associated with the local activation of different brain regions when compared to learning from an asocial CS.

### Brain functional networks associated with social and asocial classic conditioning in zebrafish

Using the correlation matrices of activity levels across the studied brain regions (i.e. matrices of co-activation), we constructed weighted networks (see methods below for details) describing the patterns of regional coactivations for each of the four combinations of experimental conditions: asocial learning (AL); social learning (SL); asocial control (AC); and social control (SC).

First, we studied how regions differently integrate during social (S) and asocial (A) learning by analyzing changes in their local network neighborhoods. This was achieved by considering the difference matrices between the learning and the control treatments in response to exposure to the social and the asocial conditions. More precisely, we computed the matrix Δ^S^ = A^S,L^ − A^S,C^, which quantifies the changes between learning and control in the social condition, and Δ^A^ = A^A,L^ − A^A,C^, which instead quantifies changes between learning and control in the asocial condition. We found no overall significant correlation between the overall restructuring of the regional neighborhoods, quantified by module of the vector of regional differences between the two conditions (i.e. the rows ∣Δ^S^_i_∣ and ∣Δ^A^_i_∣ corresponding to changes to the neighborhood of region i; Figure 3A). However, we identified a set of regions that display significant changes in activity with respect to a random null model (Figure 3B). More in detail, a significantly (≥ 95%) large amplitude of the regional changes was found for V LI_l_, Dm_r_, Dc_r_, and Ppa_r_ in social learning and for Dc_l_, A_l_, ATNI_l_, Hd_l_, Hc_l_, and VM_r_ in asocial learning, and a significantly small (< 5%) amplitude was found for OB_l_, ATNI_l_, PGZ_l_, Vl_r_, PM_r_, PPp_r_, Hav_r_, VM_r_, Had_r_, Cil_r_, NLV_r_, for social and for D_l_, Vs_l_, CP_l_, D_r_, Vc_r_, Vs_r_, A_r_, Had_r_ in asocial learning. The significance was constructed by repeatedly sampling N values uniformly at random from the Δ matrices to form a principled expectation for the amplitudes (Figure 3B). We consider then the changes in the structure of the regional neighbourhoods by computing the cosine similarity χ values between the rows corresponding to the same region of the Δ^S^ and Δ^A^ matrices (Figure 3C). We find that the majority of regions do not display significant changes with respect to a randomized version of the neighbourhood networks (see Methods). However, a few regions do display statistically significant changes. In particular, we find that regions Vl_l_ and VLI_l_ are significantly (p < 0.05, z < −1.5) dissimilar between social and asocial learning conditions. Another set of regions instead displays significant similarity between the two conditions (p < 0.05, z > 1.5): Vd_l_, Vv_l_, Dd_l_, A_l_, LH_l_, DTN_l_, GC_l_, Vd_r_, Dl_r_, ATNI_r_, TPp_r_, CP_r_, DTN_r_, GC_r_, implying that their local integration structure is conserved between conditions.

**Figure 3.**
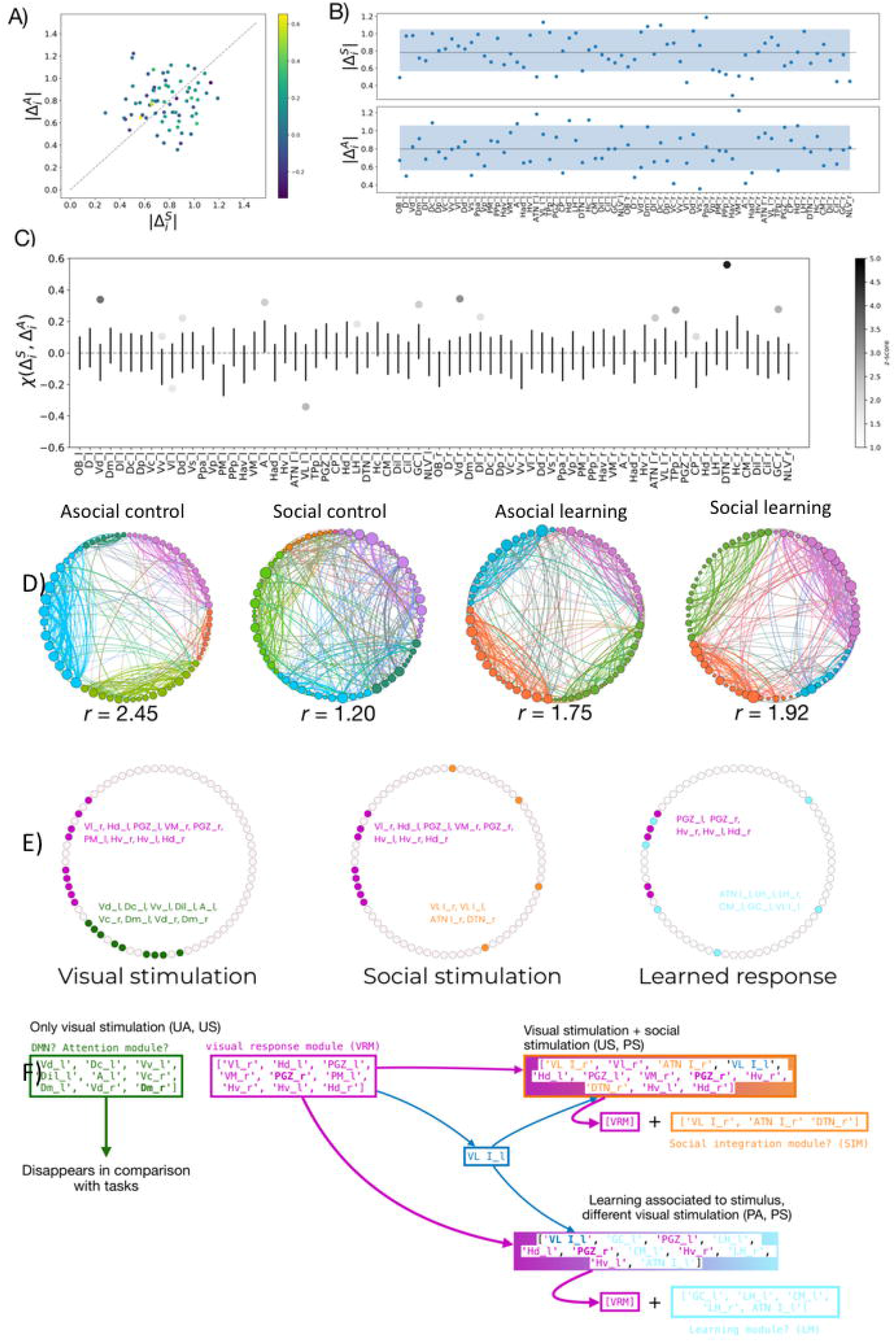
Brain networks associated with social and asocial learning. (A) Similarity of region neighborhoods between social |Δ^S^_i_| and asocial learning |Δ^A^_i_|, where the color of each data point identifies X(Δ^S^_i_, Δ^A^_i_). (B) |Δ^S^_i_| and |Δ^A^_i_| for all brain regions; the blue color band identifies the interval between the 5^th^ and 95^th^ percentiles of the randomized distribution; For details on the statistical significance see Tables S1 and S2. (C) Similarity _X_(Δ^S^_i_, Δ^A^_i_) values for all brain regions. Error bars represent one standard deviation over and below the null mean value; regions marked with dots are those with ξ values significantly different from random; the color encodes the z-score of the region’s = ξ value with respect to the random null model; For details on the statistical significance see Table S3. (D) Detection of robust functional modules in the brain networks for each treatment; within each treatment, network nodes were ordered and colored according to the module they belong to, and the degree of integration (lower r) or segregation (higher r) of the networks is provided; in all cases the measured r values are significantly larger than expected (for statistical details see Fig. S2 and Table S4), and there are differences in integration across treatments (for statistical details see Fig. S3 and Table S5). (E) Conserved brain network submodules between treatments reveal a default mode network (green module), a visual response module (purple module) a social integration module (orange module) and a learning module (blue module); regions indicated in bold font are those highlighted by the analysis based on their egonetworks. (F) Schematic representation of how the identified modules are recruited to the different tasks: the common modules present in the non-learning (i.e. unpaired US-CS) treatments (i.e. UA, US) are interpreted as a default mode network (green module) and a network responding to visual stimulation (purple module), given that the latter is also present across all four conditions; the common modules between unpaired and paired US-CS (i.e. US, PS) treatments are interpreted as a social module (blue module), since the commonality between these two treatments is the presence of a social stimulus; the common module between the two learning treatments (i.e. PA, PS) is interpreted as a learning module (blue module), since the commonality between these two treatments is the US-CS pairing during the training trials.

After having identified localized regional differences across treatments, we asked whether the network structure differs at intermediate scales (mesoscales) between treatments. This kind of deviation would signal that different information integration and elaboration strategies are used for different tasks. To this aim, we detected the modular structure of functional networks for each treatment using state of the art community detection techniques (see Figure 3D for the results of the community detection, and Methods for the robustness analysis). The first question that we can ask is whether the modular structure differs across treatments. For example, the two control (C) treatments are characterized by a slightly larger number of communities (5), with respect to the learning (L) treatments (4). It is more informative however to investigate to what degree the various communities are tightly linked within themselves with respect to with each other. To quantitatively characterize the differences among partitions, we measured the ratio r (see Methods) of the total edge weight within a community to the total weight the edges between communities. If r ≥ 1, it means that communities are denser than the inter-community medium, suggesting stronger segregation of activity within the communities as opposed to the across them. We find that the treatment asocial control (AC) has the highest r (r = 2.45), followed by the social learning (SL) (r = 1.92), and the asocial learning (AL) (r=1.75) treatments, highlighting the presence of better defined communities and tighter segregation of the functional activity within them in these treatments (Figure 3D). In contrast, the social control (SC) treatment displays a lower r value (r = 1.2), indicative of higher integration across communities rather than segregation within them (Figure 3D).

In addition to the overall balance between integration and segregation, we can ask whether these functional communities --or parts of them-- are conserved across different treatments. We did this by comparing communities between partitions corresponding to different treatments and looking for intersections between them (see Methods). We first compared the two control treatments (AC and SC). We found two conserved submodules (intersections of communities): one comprising Vd_l_, Dc_l_, Vv_l_, Dil_l_, A_l_, Vc_r_, Dm_l_, Vd_r_, Dm_r_ (green module in Figure 3 E), and a second one containing Vl_r_, Hd_l_, PGZ_l_, VM_r_, PGZ_r_, PM_l_, Hv_r_, Hv_l_, Hd_r_ (purple module in Figure 3 E). Interestingly, this second submodule appears with some modifications also when comparing the social and the learning treatments (SC, SL and AL, SL). Thus, we considered it to represent a visual response module. Then, we compared the two social treatments (SC and SL) to detect conserved submodules involved in social information processing. In this case, we found a single large conserved submodule, containing the regions VL_r_, Vl_r_, ATN_r_, VL_l_, Hd_l_, PGZ_l_, VM_r_, PGZ_r_, Hv_r_, DTN_r_, Hv_l_, and Hd_r_. Note that this module contains a large part of the second module conserved between the control treatments (AC and SC), with the addition of regions VL_r_, ATN_r_, VL_l_ and DTN_r_ (i.e. social integration module = orange module in Figure 3E). Finally, we compared the two learning treatments (AL and SL) to detect conserved submodules involved in general learning and found a single large conserved submodule containing regions VL_l_, GC_l_, PGZ_l_, LH_l_, Hd_l_, PGZ_r_, CM_l_, Hv_r_, LH_r_, Hv_l_, and ATN_l_. Note again that this submodule contains a large fraction of the second module found in the comparison between controls (AC-SC), with the additional regions GC_l_, LH_l_, CM_l_, LH_r_, and ATN_l_ (i.e. general learning module = blue module in Figure 3E). In all cases, we found that the conserved submodules include region VL_l_.

## Discussion

In this study zebrafish learned equally well a CS-US pairing using either social or an asocial CS, as there were no significant differences between the social and asocial learning treatments nor in learning acquisition during training phase neither in recall during the probe test. Importantly, we confirmed that zebrafish were able to discriminate between the social and asocial cues (i.e. CSs) used in this study and, as previously described, that they have a preference for the social cue (27). Therefore, social and asocial classic associative learning seem to be equally efficient.

When we analysed the levels of brain activation, quantified by *c-fos* expression, we found that social learning is associated with increased activity in the olfactory bulbs (OB), the ventral zone of ventral telencephalic area (Vv), the ventral habenular nuclei (Hav) and the ventromedial thalamic nuclei (VM), whereas asocial learning is associated with a decrease in activity in the dorsal habenular nuclei (Had) and in the anterior tubercular nucleus (ATN). Interestingly, all the brain regions associated with social learning have been previously implicated in learning tasks. The OB has been described as an important brain region for social learning. The cryptic cells (a subtype of cells in olfactory bulbs) are recruited in kin recognition (28), and an increase of GABA and glutamate in mitral cells is observed after training in social transmission of food preference (29–32). In our learning paradigm the CS is a visual cue and the US can be perceived either by visual or chemical cues. Thus, the increased activity of the OB cannot be explained as a direct response to the visual CS in the probe test phase, but rather as a conditioned response to it after successful pairing of the chemical US with the CS during the training phase. Thus, the expectation of food is apparently increasing the activity of the OB in anticipation of a feeding event, suggesting a modulation of olfactory perception by the social CS. The association of Vv with social learning in zebrafish is not surprising given that the lateral septum, which is its putative homologue in mammalian brains, has been implicated in several learning processes, such as auditory fear learning (33,34), contextual learning (33,35), and working memory (36). Other studies revealed the role of Vv in the processing of social information, such as social orientation (37), audience effects (38) and social exploration (39). Together, this evidence is congruent with our findings, where Vv is crucial to learning related from social cues. The lateral habenula (LHb), which is the mammalian putative homologue of the Hav (40), has also been implicated in learning and memory. For instance, inhibition of the LHb leds to deficits in spatial memory (41), object recognition (42), spatial working memory (43), aversive conditioning to cocaine (44), and complex conditioning task (45). Moreover, social behavior is also regulated by the LHb as evidenced by the decrease of *c-fos* expression in the LHb in social isolation, by the reduction in *c-fos* expression during social play (46), and by the impair of social behaviors by the activation of LHb or pre-frontal cortex (PFC) neurons and PFC-LHb projections (47–50). Finally, the VM, which is considered as a thalamic nuclei in zebrafish (51), belongs to the cortico-basal ganglia-thalamic loop circuit in mammals, in which the basal ganglia receive inputs from the cortex and transfers them back to frontal and motor cortex via the VM (52). This circuitry is also connected with LHb allowing animals to adjust the salience and valence of stimuli.

In contrast, asocial learning in zebrafish is associated with other brain regions, namely the dorsal habenular nuclei (dHb) and the anterior tubercular nucleus (ATN). The dHb, which is homologous of the medial habenula in mammals (mHb), receive inputs mainly from the limbic system, and sends outputs to the interpeduncular nucleus, which in turn regulates activity dopamine (DA) and serotonin (5HT) neurons (53–55). Evidence in both mice and zebrafish supports our results that suggest dHb to be related to asocial learning. Ablation of mHb induces deficits in long-term spatial memory (54), complex learning paradigms (54) and fear learning (56,57). In contrast, our findings reveal a decreased expression of *c-fos* associated with asocial learning, probably due to a disinhibitory mechanism. The ATN is homologous of the ventromedial hypothalamus (VMH) in mammals, a brain region that has also been related to learning processes with strong *c-fos* expression after fear conditioning (58) and recall of conditioned fear (59). The role of VMH in learning processes in mammals can be explained by the afferents from the amygdala (BLA and MEA), a brain region clearly shown to be involved in learning processes (58). In summary, social and asocial learning in zebrafish are associated with changes in activity in different sets of brain regions known to be involved in learning in other species.

We have also studied the structure of brain networks in relation to the two types of learning. The regional similarity data also reveals a lack of correlation between social and asocial learning, and a large amplitude of the change in the neighborhood structure(i.e. implying more marked local changes in network structure) for different regions in the two types of learning (i.e. social: VLI_l_, Dm_r_, Dc_r_, Ppa_r_; asocial: Dc_l_, A_t_, ATNI_l_, Hd_l_, Hc_l_, VMr). Furthermore, the community detection results revealed a robust modularity of the brain networks across all treatments, with the social learning treatment displaying a significantly higher integration across communities than the asocial one. Finally, we looked into the composition of shared submodules in an attempt to frame and contextualize them. We offer a hypothesis on the basis of these results. The treatments AC-SC share the visual response to a stimulus but include no learning on the part of the individuals. We can imagine therefore that the shared submodules will encode the simple reaction of the animal to the appearance of a visual stimulus carrying no semantic meaning (as the animal has not learned to associate it with food). The two submodules should therefore code for the generalized attention (AM) and the visual response mechanisms (VRM) (Fig. 3 F). In the comparison SC-SL, the commonality lies in the presence of a social visual stimulus. We would expect therefore to see a recruitment of the VRM with a potential additional recruitment of other regions responsible for social recognition. From this perspective, the single conserved submodule that we found in the SC-SL supports this interpretation, being largely composed by the VRM regions and a few additional ones, that we now denote as social integration module (SIM) (Fig. 3 F). Along the same line, the AL-SL should highlight the submodule specific to learning and reacting to the food stimulus, independently from the type of visual stimulus. We find again a single large conserved submodule, that includes about 50% of the VRM regions plus a new set of regions with no overlap with the SIM. Arguably, these additional regions should be responsible for the learned response and association with the food stimulus, and we denote them as the learning module (LM) (Fig. 3 F). Finally, the region VL_l_ constitutes a glaring exception, as it appears in all the submodules that we described so far. This general presence might suggest that it has a generic role in information integration across different areas.

In summary, here we show that social and asocial learning are associated with localized differences in brain activity that are paralleled by the segregation of brain modules that seem to serve subsets of cognitive functions, such as a visual response module, a social integration module and a learning module that is shared between the two types of learning. Together, our results provide the first experimental evidence for the occurrence of a general-purpose learning module that is apparently modulated by different patterns of localized activity in social and asocial learning.

## Methods

### Animals

Zebrafish (*Danio rerio*) were 5 months old wild-type (Tuebingen strain) males, bred and held at Instituto Gulbenkian de Ciência Fish Facility (Oeiras, Portugal). Fish were kept in mixed-sex groups, at 28ºC, 750 μs, pH 7.0 pH in a 14L:10D photoperiod and fed twice a day (except on the day of the experiments) with freshly hatched *Artemia salina* and commercial food flakes.

### Ethics statement

All experiments were performed in accordance with the relevant guidelines and regulations, reviewed by the Instituto Gulbenkian de Ciência Ethics committee, and approved by the competent Portuguese authority (Direção Geral de Alimentação e Veterinária, permit number 0421/000/000/2017).

### Behavioral paradigm

One day before the experiment, fish were moved to the home tanks (1.5L, 12.5 cm x 12.5 cm x 12.5 cm) where they only had visual and chemical access to a mix shoal of 4 animals (2 familiar males and 2 familiar females).

The experiment was subdivided into three phases: acclimatization, training and probe test. In the acclimatization phase, after one minute in the start box, animals were allowed to swim freely in the tank for 9 min, during which, they were attracted to all arms of the plus-maze with bloodworms, so that they became familiar with the whole maze. In the training phase, animals were trained in daily sessions of trials per session for 6 days. In the paired groups, animals had the CS and the US presented together in the same arm, and received a reward (bloodworm) when this arm was chosen (that changed on each trial in a pseudo-randomized way, within and between individuals); when another arm was chosen, animals stayed one minute in the chosen arm, and then they were conducted to the right arm, using a hand net where they receive the reward. In the unpaired groups, the animal spent 2 minutes in the chosen arm since the CS and US were never presented together. In both groups, when individuals reach the RoIof the chosen arm the start box was closed to avoid the animal change its decision. In the social treatments, the CS stimulus presented at the end of the arm was a static, 2D photography of a zebrafish. In the asocial treatments a digitally drawn circle with the same visual target area and the same mean color of the zebrafish-stimulus was used. After each training session, individuals returned to their home tank.

In the probe test (24h after the last training trial), animals were only exposed to the CS for 2 min. The CS was then removed and the animal remained in the tank for 30 min to achieve the peak of expression for *c-fos* (60).

A preference test was performed to assess if individuals prefer social to asocial CS’s. For this purpose, we used a rectangular tank (5L, 30cm x 15 cm x 15 cm) with the stimuli presented on each side (e.g. social stimulus in the left and asocial stimulus in the right side, in a randomized way between individuals). Individuals were placed in a start box for 2 min with transparent partition, and the time spent in both RoI’s was compared.

To demonstrate that individuals can discriminate between the two CS used (i.e. social and asocial CSs), we performed a discrimination task. In this case, we trained fish (one-minute trial, 8 trials/day for 5 days) to associated one CS to a reward (food) and the other to a punishment (netting) (e.g. social stimulus in half of the animals was associated with food and in the other half it was associated with threat). In the probe test, only the CSs were presented, and we measured the duration spent by the focal fish in each arm; if individuals were able to discriminate between the two stimuli, they should prefer the arm associated with reward independently of their initial preference for the social stimulus.

In all experiments, the behavior was recorded with a digital camera for subsequent analysis using a commercial video tracking software (EthoVisionXT 8.0, Noldus Inc. the Netherlands).

### Brain collection

Animals were sacrificed with an overdose of Tricaine solution (MS222, Pharmaq; 500–1000 mg/L) and sectioning of the spinal cord. The brain was macrodissected under a stereoscope (Zeiss; Stemi 2000) and immediately collected to 4% PFA solution in 0.1M PB and kept overnight at 4º C. After cryopreservation (34% sucrose in 0.1M PB ON at 4ºC), the brains were embedded in mounting media (OCT, Tissue teck) and rapidly frozen on liquid nitrogen. The coronal sectioning was performed on a cryostat (Leica, CM 3050S) at 16 um, sections were collected onto SuperFrost glass slides and stored at -20ºC.

### In situ hybridization for the immediate early gene *c-fos*

Chromogenic RNA in situ hybridization (CISH) was carried out according to a standard protocol available upon request from the lab of Professor Marysia Placzek, University of Sheffield, briefly described below. For the generation of *c-fos* probes, a pBK-CMV vector containing the *c-fos* cDNA (Genebank: CF943701) was cut with the restriction enzyme BamHI (antisense) and EcoRI (sense) to generate templates for in vitro transcription. Digoxigenin-labeled *c-fos* sense and antisense probes (11277073910, Merk (Roche), UK) were then synthesized through in vitro transcription of 1 mg template with T7 polymerases (M0251, New England Biolabs). The sections were fixated in 4% PFA, washed in PBS, rinsed in 0.25% acetic anhydride in 0.1 M tri-ethanolamine for 10 min and washed 3 times in phosphate buffer saline (PBS). An incubation in pre-hybridization buffer (hybridization solution without yeast RNA, minimum 3 hours) was done in order to prepare the tissue for receiving the probe diluted in hybridization solution (probe dilution: 1:40 ∼4 ng/ul final concentration). The hybridization buffer contained 50% formamide, 5 x SSC (pH 7.0), 2% blocking powder, 0.1% triton X-100, 0.5% CHAPS, 1 mg/ml yeast RNA, 5mM EDTA and 50 ug/ml heparin. The hybridization incubation was performed at 68°C for 24 h. Following hybridization, the sections were treated with secondary antibody anti-dig-ap (1:1000, 11093274910, Merk (Roche), UK), after a series of several washes decreasing concentrations of SSC, until 0.1× SSC. The tissue was then mounted onto GlicerolGel (GG1, Merk) coated slides and left to air dry.

### Cell Counting

The slides were imaged using a tissue scanner (NanoZoomer Digital Pathology, Hamamatsu). A whole brain screening was performed to select the brain nuclei with higher *c-fos* activity to be counted (see list in Table1). The areas were manually drawn and the signal automatically quantified using the Icy software (created by the Quantitative image analysis unit at Institut Pasteur). The sum of the cross sections was used as an individual measure to each area side.

### Statistical Analysis

A N-1 chi-square test for proportions was used to compare the proportions of learners, non-learners and non-retention individuals, relative to the total amount of individuals in each treatment. Non-parametric linear regressions were performed to compare the learning curves across the 4 experimental treatments. To assess differences between the experimental treatments a non-parametric test with the location on the plus maze (target arm, front, left or right arms) as within factor and social (social, asocial) and learning (learners, non-learners) factors as between factors, was used. Planned comparisons followed by Benjamini and Hochberg’s method for p-value adjustment were also used to assess the brain regions associated with social learning (social learning vs social control) and asocial learning (asocial learning vs asocial control).

The effect of social learning on brain activity in the probe test was assessed by a non-parametric test with laterality (number of *c-fos* positive cells on each nuclei on the left and right side of each brain) as repeated measure and social and learning as between factors area-by-area (OB, D, Vd, Vc, Vv, Vl, Dc, Dl, Dm, Dp, Vs, Dd, PPa, Vp, PM, PPp, Had, Hav, A, VM, VL, Hv, ATN, LH, Hd, CP, TPp, PGZ, Hc, DIL, CIL, DTN, NLV GC, CM). Planned comparisons followed by Benjamini and Hochberg’s method for p-value adjustment were used to assess the brain areas associated with social learning (social learning vs social control) and asocial learning (asocial learning vs asocial control).

### Network analysis

#### Construction of the correlation graph tower

Functional networks for brain connectivity are usually built starting from vectorial information on the individual regions. In network neuroscience, this means computing a similarity measure (e.g. Pearson correlation) between the timeseries of brain parcels in fMRI, or channels in EEG. In the case of social learning experiments the correlations of number of positive cells for each pair of brain nuclei, for within each experimental treatment was computed, however we have access to a single measure per region for each specimen. In addition, the number of specimens is typically limited. In turn this makes the estimation of the actual correlation (or similarity) between regions more complicated.

To account for this, we take inspiration from standard bootstrapping and, instead of defining a single network, we construct a set of possible networks leaving out some of the specimens information’s.

More precisely, consider the case of *M* specimens, each with one sample reading *x*_*i*_ for each of the N brain regions. Given a similarity metric *ω*, typically one would consider the *M*-dimensional vectors *x*_*i*_, where *i* labels the regions, and then compute the similarities *ω*_*ij*_ = *ω*(*x*_*i*_, *x*_*j*_) for all pairs *ij*. The resulting weighted matrix *Ω* is then interpreted as the adjacency matrix of a functional network.

For a given m < *M*, we will instead consider all the 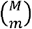 combinations {*γ*} of *m* specimens and compute the corresponding functional graph *Ω*_*γ*_. We will refer to the collection of graphs obtained in this way as a *graph tower Ω*_*Γ*_, where each of the combinations can be considered as a *graph layer*. Similar constructions are used for multilayer and multiplex networks with the notable difference that graph towers do not have edges connecting the different layers.

The advantage of this construction is that each layer in the graph tower represents a different instance of the network bootstrapping. In this way, observables computable on a single layer can be bootstrapped across multiple ones. This construction has naturally one parameter, the sampling number m, which needs to be chosen on the basis of data-driven considerations or the robustness of the resulting networks.

#### Selection of threshold

Correlation networks are usually fully connected weighted networks. It is however common to sparsify them by retaining only edges that have a weight larger than a certain weight threshold. Another common practice is to choose a target density ρ for the graph and add edges to the network starting from the strongest ones until the density is reached. Given a graph (layer) Ωγ, we will denote the graph obtained using a threshold Ωγ at density ρ as Ω^ρ^_γ_. While the sparsification is often required to highlight the network properties of the system and to filter out weaker correlations, there is no commonly accepted method to choose such thresholds. Typically, the adopted methods depend strongly on the specific application and are developed ad-hoc. Overall, most existing methods rely either on considerations on the data used to construct the correlation matrix (e.g. the timeseries in neuroimaging), or on the local structure of the network (e.g. disparity filter).

Here, we take a different route and leverage the graph tower structure to choose the threshold value. We will work using the density as threshold, but the same argument can be replicated using weights in a straight-forward manner. For each edge ij, we can consider the set of edge weights {ω_ij_}_γ_ across all layers {γ}. Denoting respectively μ(ω_ij_) and σ(ω_ij_) as the mean and standard deviation of the ω_ij_ over the layers, we can associate to Ω a mean heterogeneity

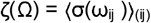

and a mean coefficient of variation

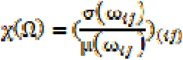

Denoting as Ω^ρ^_Γ_ the graph tower thresholded at density ρ, the two quantities above can be computed as a function of the threshold density ρ. In Figures S1A-B we report the dependence of ζ and χ. We find a clear change in the heterogeneity patterns at around ρ_0_ = 0.05. In particular, near ρ_0_ the mean heterogeneity ζ is still minimal, while the coefficient of variation χ peaks before starting to decrease again. In Figure S1C we report for comparison a recent method for density thresholding proposed in (61). This method identifies ρ_0_ as the density that maximizes the quantity 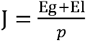, where Eg and El are respectively the global and local network efficiency (62). Interestingly, we find the threshold density identified by J is very close to the ρ_0_ identified by ζ and χ, suggesting that our construction based on the heterogeneity patterns of Ω_Γ_ captures a critical point in network structure. In the rest of the paper, all the networks will be thresholded at this density ρ_0_.

#### Comparison of region connectivity profiles

We are interested in comparing how brain regions is linked to each other in the various conditions and tasks. During different tasks, the same brain region might be performing different functions by changing how it links to the other regions, its local context (also called egonetwork). Given two conditions A and B with associated matrices A^A^ and A^B^, we can measure then the change in the local environment of a region i by computing the amplitude and direction of the change. The amplitude of the change can be quantified by considering the norm of the difference between the row vectors associated to i in the two conditions A_i_^A^ = (A_i0_, … A_iN−1_) and A_i_^B^ = (A ^A^ _i0_, … A ^B^ _iN−1_). That is, we calculated the vector difference between the two rows Δ^AB^_i_ = ∣ A^A^_i_ − A^B^_i_ ∣ and then take its norm ∣Δ^AB^_i_ ∣. To quantify the direction of the change, we can instead the cosine similarity between A^A^_i_ and A^B^_i_ :

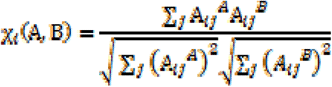

#### Detection of robust functional modules

Communities were computed using the Leiden community detection method (63) on the graph tower matrices, averaged over all the bootstrapping samples at fixed density *p* = 0.07. To increase the robustness of the detection, for each treatment, we repeated the community detection 100 times. From the 100 candidates partitions we extracted the central partition as described in (64) and associated the resulting partition to the treatment under analysis.

To quantitatively characterize differences among partitions, we measure the ratio *r* of total edge weight within a community with that of the edges between communities. More specifically, for partition P with m communities we compute the (*m* x *m*) matrix P, defined as:

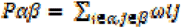

Where α, β = 0, … *m*-1label the modules of *P*, and *ωij* is the edge weight between regions I and j. We then compute the ratio of average intra-community edge weights as follows:

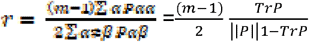

Which measures the ratio of the average weight on the diagonal of P_α, β_ to the average off-diagonal weight.

We would also like to identify modules, or parts of modules, that are shared across partitions corresponding to different treatments. One method to quantify this is to study the overlap between pairs of modules: consider partitions Px = {C^0^_x_, C^1^_x_ … C^m^_x_} and P_y_ == {C^0^_y_, C^1^_y_ …C^l^_y_} for treatments x and y; for each pair of modules (C^i^_x_, C^j^_y_), we compute the intersection J^ij^_x,y_ = C^i^_x_ ∩ C^j^_y_. To establish significance, we employ a permutation test based on a null distribution for the size of intersections p_0_(∣J∣): for each pair (C^i^_x_, C^j^_x_), we sample uniformly at random 10000 pairs of nodesets with cardinality respectively ∣C^i^_x_ ∣ and ∣C^j^_x_ ∣ and compute the size of their intersection ∣J∣. We then retain the submodule J^i,j^_x,y_ iff ∣J^i,j^_x,y_∣ ≥ μ(∣J∣) + 3σ(∣J∣), where μ(∣J∣) and σ(∣J∣) are the first two moments of p_0_(∣J∣).

## Acknowledgements

This study was funded by a research grant from Fundação para a Ciência e a Tecnologia (FCT, Portugal, grant reference: PTDC/BIA-ANM/0810/2014) awarded to RFO. JP was supported by a FCT doctoral fellowship (PTDC/BIA-ANM/0810/2014). We would like to acknowledge all members of Dr. Vincent Cunliffe laboratory, in particular Dr. Helen Eachus for technical support, and Dr. Catarina M. Henriques for technical support in the *in situ* hybridization in brain slices of zebrafish.

