## Supplemental material for "Social and asocial learning in zebrafish are encoded by a shared brain network that is differentially modulated by local activation"

**This PDF file includes:**

Tables S1 to S5

Figs. S1 to S3

**Supplementary Tables**

**Table S1.** P-values for regions with significantly similar neighbourhoods between treatments S and A.

| Region | S | A |
| --- | --- | --- |
| VL I_l | 0.0241 | - |
| Dm_r | 0.0379 | - |
| Dc_r | 0.0337 | - |
| Ppa_r | 0.0132 | - |
| Dc_l | - | 0.0365 |
| A_l | - | 0.0408 |
| ATN I_l | - | 0.0092 |
| Hd_l | - | 0.027 |
| Hc_l | - | 0.0242 |
| VM_r | - | 0.0049 |

**Table S2.** P-values for regions with significantly dissimilar neighborhoods between treatments S and A.

|  | S | A |
| --- | --- | --- |
| OB_l | 0.0096 | - |
| ATN I_l | 0.0123 | - |
| PGZ_l | 0.0136 | - |
| Vl_r | 0.0018 | - |
| PPp_r | 0.0247 | - |
| Hav_r | 0.0 | - |
| VM_r | 0.0163 | - |
| Had_r | 0.007 | 0.0283 |
| Cil_r | 0.0027 | - |
| NLV_r | 0.003 | - |
| D_l | - | 0.013 |
| Vs_l | - | 0.016 |
| CP_l | - | 0.0272 |
| D_r | - | 0.0089 |
| Vc_r | - | 0.0008 |
| Vs_r | - | 0.0001 |
| A_r | - | 0.0011 |
| TPp_r | - | 0.0477 |

**Table S3.** P-values and z-scores for significant similarity values χ($\Delta$^S^_i_,$\Delta$^A^_i_). Negative z-scores correspond to regions with significantly smaller similarity between S and A, implying that their neighborhood changes strongly. Positive z-scores correspond to regions whose neighborhood is significantly conserved between S and A.

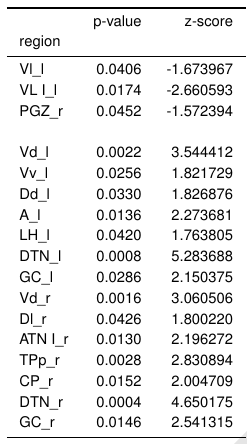

**Table S4.**$r$ values with significance.

| treatment | $r$ | p-value |
| --- | --- | --- |
| (P, A) | 1.750281 | 1.0000 |
| (P, S) | 1.926310 | 1.0000 |
| (U, A) | 2.522384 | 1.0000 |
| (U, S) | 1.203354 | 0.9995 |

**Table S5.**$\Delta r$ values with significance.

| treat 1 | treat 2 | $\Delta r=r_{1}$ - $r_{2}$ | p-value |
| --- | --- | --- | --- |
| (P, A) | (P, S) | -0.176029 | 0.0485 |
| (P, A) | (U, A) | -0.701090 | 0.0000 |
| (P, A) | (U, S) | 0.656638 | 1.0000 |
| (P, S) | (U, A) | -0.525061 | 0.0000 |
| (P, S) | (U, S) | 0.832667 | 1.0000 |
| (U, A) | (U, S) | 1.357728 | 1.0000 |

**Supplementary Figures**

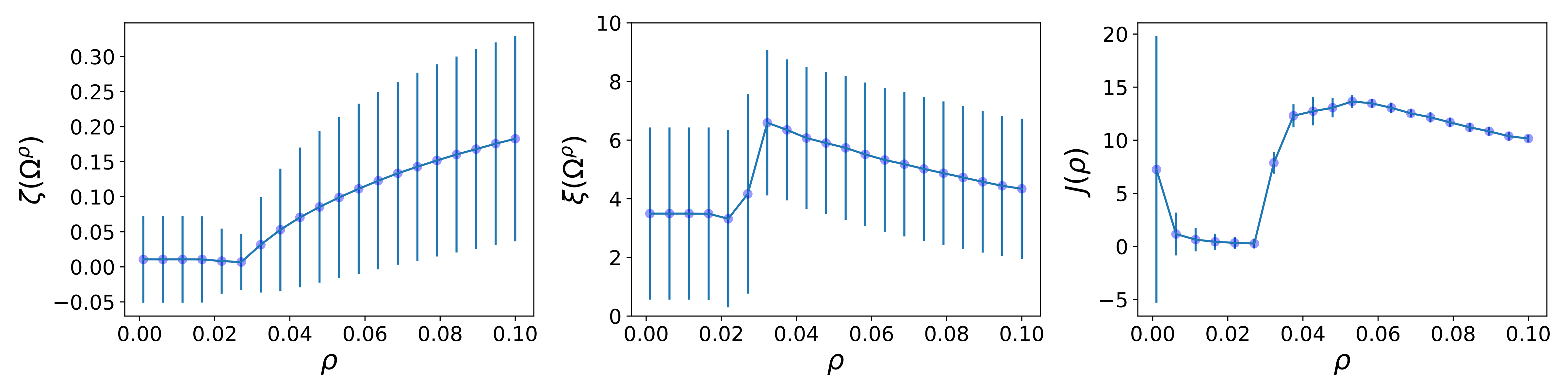

**Figure S1.** Heterogeneity and efficiency of graph tower as function of density ρ.

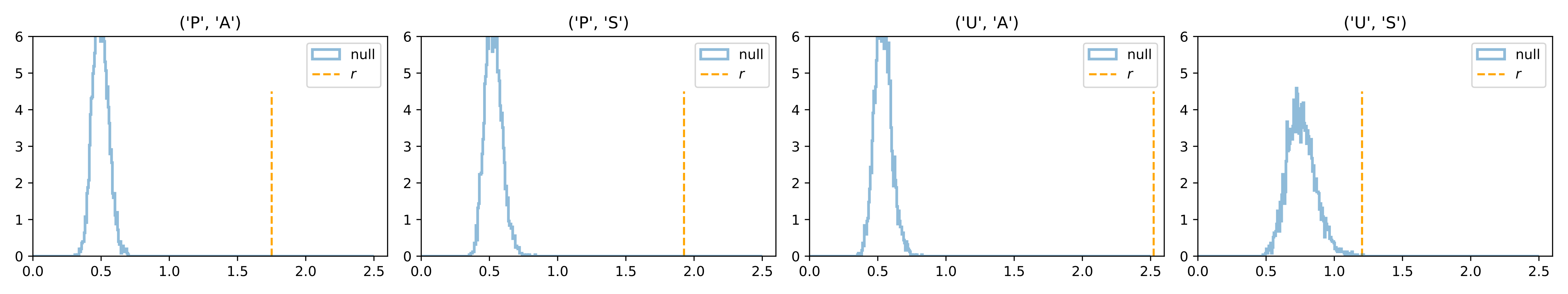

**Figure S2.** Null distributions for r values. The panels show the null distributions p(r') (blue) and the r values obtained from data for each treatment. In all cases, the measured r are (very significantly) larger than expected from the null model.

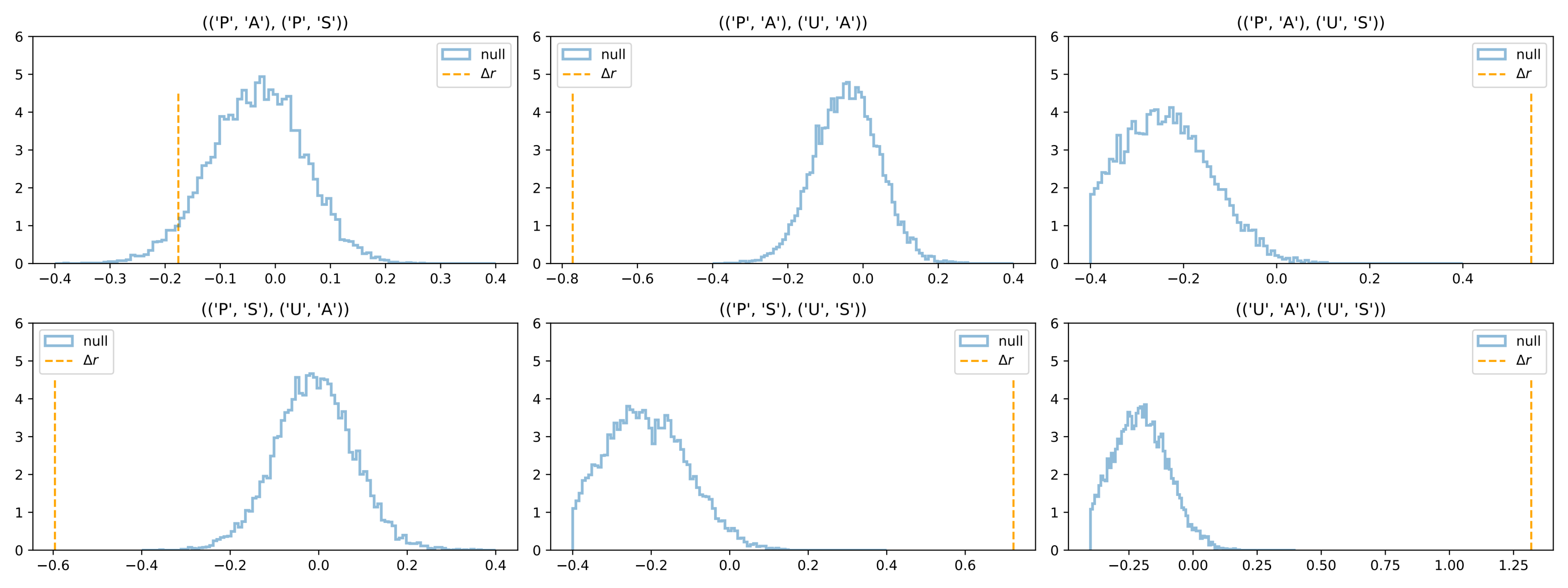

**Figure S3.** Null distributions for Δr values. The panels show the null distributions p(Δr') (blue) and the Δr values obtained from data for each pair of treatments. The measured Δr are very significantly different from the null expectation, with the exception of the comparison PA-PS which marginally significant for the significance threshold α=0.05.
